# White Matter Slow-4 fALFF as a Complementary Marker in the Multimodal Alzheimer’s Disease Biomarker Landscape

**DOI:** 10.64898/2026.09.17.752468

**Authors:** Yali Huang, Arthur W. Toga, Lu Zhao, the Alzheimer’s Disease Neuroimaging Initiative

## Abstract

**INTRODUCTION:** White matter (WM) functional activity, quantified as slow-4 fractional amplitude of low-frequency fluctuations (fALFF; 0.027–0.073 Hz), may reflect Alzheimer’s disease (AD) pathology, but its utility relative to established structural imaging markers remains poorly characterized.

**METHODS:** We analyzed 369 ADNI-3 participants with baseline resting-state fMRI. Global WM slow-4 fALFF (JHU-20 atlas) was compared against hippocampal volume, entorhinal cortical thickness, FA, and MD across 11 outcomes: plasma biomarkers (pTau217, Aβ42/40, NfL, GFAP; n = 108), PET (amyloid Centiloid, tau SUVR, FDG; n = 66–207), and cognition (MEM, EF, mPACC, MMSE; n ≈ 222–225) using head-to-head benchmarking and variance decomposition.

**RESULTS:** WM fALFF was significantly associated with GFAP (partial RZ = 5.35%), amyloid PET (partial RZ = 2.17%), tau PET (partial RZ = 3.52%), executive function (partial RZ = 2.61%), and mPACC (partial RZ = 3.53%). In trimmed models, fALFF explained unique variance in NfL and GFAP beyond FA and MD (unique ΔRZ: 3.65% and 4.45%) — nearly 8-fold greater than DTI’s unique contribution for NfL. Gray matter markers showed larger associations with cognition and PET pathology. The incremental value of fALFF was modality-specific: it persisted beyond DTI markers (NfL, GFAP, amyloid PET, executive function, mPACC) and beyond gray matter markers (NfL, GFAP), but in every domain — plasma included — fALFF was no longer significant when all five imaging markers were entered simultaneously.

**DISCUSSION:** WM slow-4 fALFF captures neurodegeneration-related variance — particularly astroglial reactivity and axonal injury — incremental to DTI microstructure and partially independent of gray matter atrophy. These findings position WM functional activity as a complementary, non-invasive imaging marker in the multimodal AD biomarker landscape.

**Research in Context:** *Evidence before this study:* White matter (WM) resting-state BOLD signals carry physiologically valid functional information. WM slow-4 fALFF is reduced in preclinical Alzheimer’s disease (AD) and correlates with amyloid burden and cognition. Direct benchmarking against established structural imaging markers within matched samples has remained scarce.

*Added value of this study:* In 369 ADNI-3 participants, we conducted a direct same-sample head-to-head comparison of global WM slow-4 fALFF versus hippocampal volume, entorhinal thickness, global FA, and MD across 11 pre-specified AD biomarker and cognitive outcomes. WM fALFF showed the largest covariate-adjusted association with GFAP (partial RZ = 5.35%) and uniquely explained ∼8× more NfL variance than DTI in trimmed models, contributing non-redundant functional information.

*Implications of all the available evidence:* Global WM slow-4 fALFF, derived from standard resting-state fMRI, is a non-invasive complement to structural imaging that showed associations with markers of astroglial reactivity and neuroaxonal injury, and could augment multimodal AD biomarker panels without additional scanning cost.

## 1. Introduction

Dementia affected an estimated 57 million people worldwide in 2019, a number projected to reach 153 million by 2050 as populations age, and Alzheimer’s disease (AD) is its leading cause [1]. Despite decades of research, early detection remains a central challenge: by the time clinical symptoms emerge, substantial neurodegeneration has already accumulated [2]. The development and validation of sensitive, accessible, and complementary biomarkers — capable of capturing distinct pathological dimensions across the AD continuum — is therefore a research priority of the highest clinical importance.

The revised AT(N)(I) diagnostic framework classifies AD biomarkers into four categories: amyloid-β pathology (A), tau pathology (T), neurodegeneration or neuronal injury (N), and inflammation or immune activity (I) [3,4]. Plasma Core 1 biomarkers, particularly phosphorylated tau-217 (pTau217) and the amyloid-β42/40 ratio, have demonstrated high diagnostic accuracy for AD and are now recommended for use in research and selected clinical contexts [5,6,7]. Plasma neurofilament light chain (NfL) and glial fibrillary acidic protein (GFAP) serve as non-specific N-class and I-class markers reflecting neuroaxonal injury and astroglial reactivity, respectively [4]. Structural MRI — including hippocampal volume and entorhinal cortical thickness — and PET-based amyloid and tau quantification remain central to research protocols and clinical trials [3], and combined amyloid and glucose-metabolism PET measures predict longitudinal cognitive decline in ADNI [8]. Yet each established modality has limitations: invasiveness, cost, and limited accessibility constrain PET and cerebrospinal fluid measures, while structural MRI captures atrophy that may follow rather than precede molecular pathological events. There is accordingly sustained interest in non-invasive imaging markers that can complement the established biomarker framework.

White matter (WM) involvement is an early and pervasive feature of AD pathophysiology. Diffusion tensor imaging (DTI) measures of fractional anisotropy (FA) and mean diffusivity (MD) document widespread microstructural disruption of major fiber tracts in MCI and AD dementia, and have been associated with amyloid and tau burden, neurodegeneration markers, and cognitive decline [9,10]. Beyond structural integrity, converging evidence indicates that resting-state BOLD signals recorded from WM voxels carry meaningful functional information. WM functional networks with reproducible spatial organization have been identified across independent datasets [11,12], WM BOLD waveforms synchronize with external stimuli in a fiber-tract-specific manner [13], and intracranial electrophysiological recordings have provided direct evidence that WM BOLD functional connectivity reflects neural synchrony [14]. These observations establish WM resting-state BOLD signals as physiologically interpretable indices of integrated white matter activity, distinct from and complementary to structural DTI measures.

The fractional amplitude of low-frequency fluctuations (fALFF) — defined as the ratio of power in a targeted low-frequency band to total spectral power — has been widely applied in gray matter resting-state fMRI research as a measure of spontaneous neural activity [15]. When restricted to the slow-4 frequency band (0.027–0.073 Hz), fALFF demonstrates superior test-retest reliability compared with full-band ALFF and the adjacent slow-5 band [16], and shows broader, more sensitive discrimination of preclinical AD stages relative to slow-5 [17]; frequency-dependent differences between slow-4 and slow-5 were first documented in amnestic mild cognitive impairment [18]. Importantly, a physiologically distinct spectral peak centered at approximately 0.06 Hz has been identified in WM BOLD signals — falling squarely within slow-4 — whose amplitude declines with age and has been linked to glymphatic and perivascular clearance mechanisms [19]. These convergent lines of evidence motivated the a priori selection of the slow-4 band for WM fALFF quantification in the present study.

Preliminary evidence supports the relevance of WM fALFF to AD pathology. Chang et al. demonstrated that bundle-wise WM fALFF is reduced in preclinical AD (amyloid-positive cognitively normal individuals) and associates with amyloid burden and cognitive performance [20]. Yan et al. showed that WM ALFF/fALFF abnormalities predict AD diagnosis independently of gray matter atrophy [21]. In an earlier ADNI multimodal study, white matter lesion load was associated with resting-state fMRI activity and amyloid PET, though not with FDG-PET, in MCI and early AD [22]. However, the degree to which WM slow-4 fALFF provides information unique relative to — or complementary to — established structural imaging markers (hippocampal volume, EC thickness, FA, MD) across the full spectrum of AD-relevant outcomes, including plasma biomarkers, PET, and cognitive composites, has received limited systematic evaluation. Without such head-to-head benchmarking within matched samples, the incremental and relative value of functional WM imaging remains difficult to quantify objectively.

The present study addresses this gap using data from the Alzheimer’s Disease Neuroimaging Initiative (ADNI). We quantified global WM slow-4 fALFF across 20 major JHU white matter tracts and compared its associations with 11 pre-specified AD-relevant outcomes — four plasma biomarkers (pTau217, Aβ42/40, NfL, GFAP), three PET measures (amyloid Centiloid, meta-temporal tau SUVR, FDG SROI.AD), and four cognitive composites (ADNI-MEM, ADNI-EF, mPACC, MMSE) — against four established structural imaging markers within the same complete-case samples. Variance decomposition analyses further quantified the unique contributions of fALFF and structural markers in modality-specific trimmed models. Our goal was not to establish WM fALFF as a replacement for existing biomarkers, but to systematically characterize its position within the multimodal AD biomarker landscape as a non-invasive functional complement.

## 2. Methods

### 2.1 Study Design and Participants

Data were obtained from the Alzheimer’s Disease Neuroimaging Initiative (ADNI; adni.loni.usc.edu) [23,24,25]. ADNI is a longitudinal multicenter study designed to develop and validate biomarkers for the early detection and tracking of Alzheimer’s disease, enrolling cognitively normal (CN) older adults, individuals with mild cognitive impairment (MCI), and those with AD dementia diagnosed according to National Institute on Aging–Alzheimer’s Association criteria [26], across approximately 60 North American sites. ADNI was launched in 2003 under principal investigator Michael W. Weiner, MD, with funding from the National Institute on Aging, the National Institute of Biomedical Imaging and Bioengineering, and private sector partners. All participating sites received institutional review board approval, and all participants provided written informed consent.

The present cross-sectional analysis drew on the ADNI-3 phase [25]. All resting-state fMRI scans analysed here were acquired under the ADNI-3 protocol between February 2017 and April 2023, and the sample comprised participants newly enrolled in ADNI-3 together with rollover participants originally enrolled in ADNI-1, ADNI-GO, or ADNI-2. Participants were eligible if they had a baseline resting-state fMRI scan available together with valid age and years of education. Resting-state fMRI scans with mean framewise displacement exceeding 0.5 mm were excluded to limit the influence of head motion. After these criteria, 369 participants were retained as the primary analytic cohort. Depending on the availability of each additional imaging or biomarker modality, analysis-specific complete-case subsamples were used (see Sections 2.4–2.7 and Results Section 3.1). Demographic and clinical characteristics of the analytic sample are shown in Table 1.

**Table 1.**
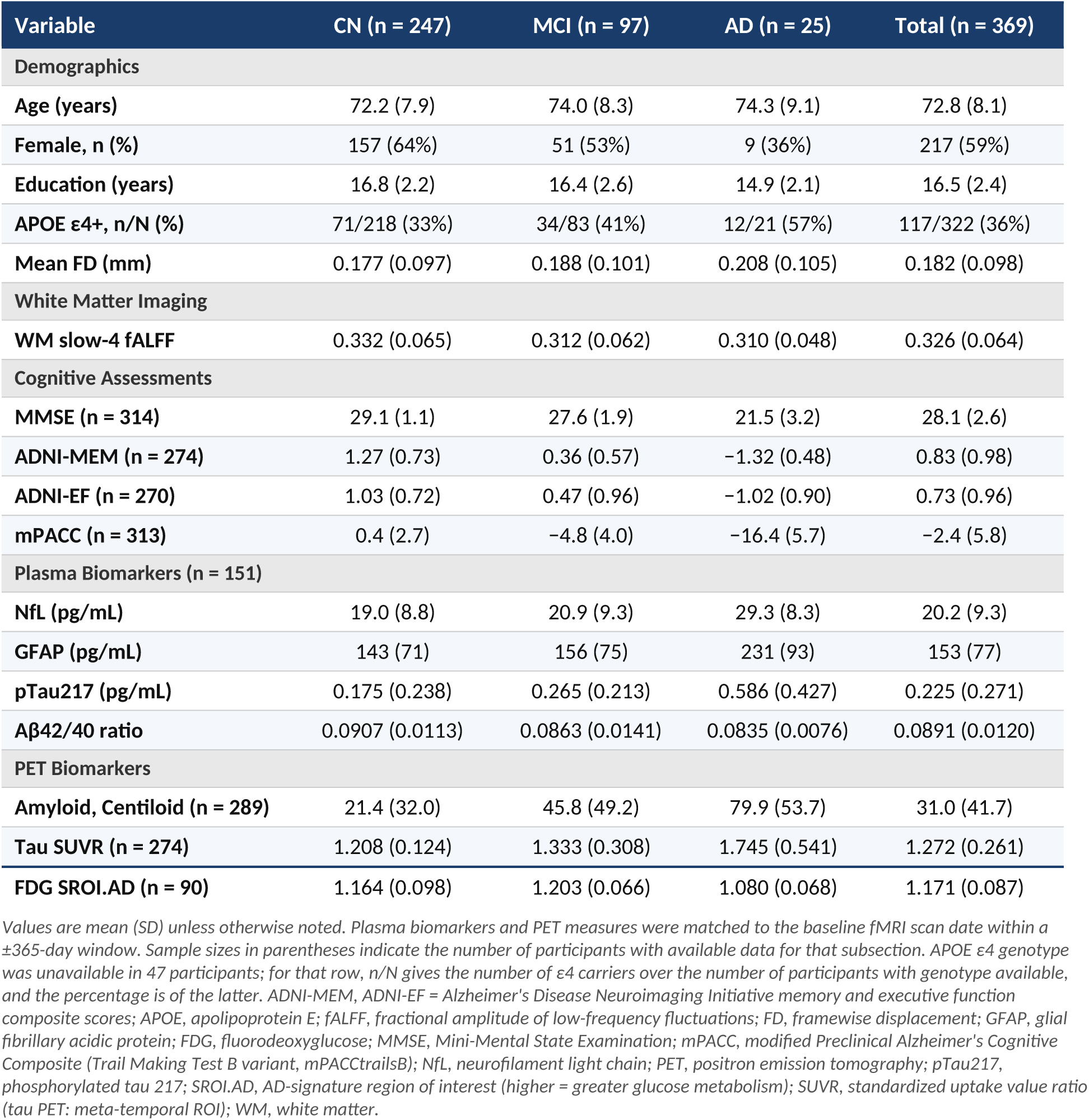
Participant characteristics by diagnostic group.

### 2.2 MRI Acquisition and Preprocessing

Resting-state fMRI was acquired with the ADNI-3 basic task-free fMRI protocol [25]: a single-echo gradient-echo echo-planar sequence acquired with eyes open (ADNI series description “Axial rsfMRI (Eyes Open)”), TR = 3,000 ms, 3.4 mm isotropic voxels, 48 axial slices, and 197 volumes (≈9.9 min). Every scan included in this study used this sequence; none of the multiband (“advanced”) ADNI-3 resting-state acquisitions were included. Resting-state fMRI data were preprocessed using fMRIPrep 25.1.4 (RRID:SCR_016216) [27,28]. Preprocessed BOLD derivatives — motion-corrected BOLD time series in MNI152NLin6Asym space, together with the accompanying confound tables — were downloaded directly from ADNI. fMRIPrep internally performed T1w bias-field correction, brain extraction, T1w-to-BOLD coregistration via ANTs, and nonlinear normalization to standard space. Analyses used the unsmoothed preprocessed BOLD series; no additional spatial smoothing was applied. Repetition time was read from each scan’s JSON sidecar rather than assumed, and was 3,000 ms for every scan in the analytic sample. The first 10 volumes of each series were discarded before spectral estimation, leaving 187 volumes (561 s) and a spectral resolution of 0.0018 Hz. Isolated high-motion volumes (FD > 5 mm) were replaced by the average of their two temporal neighbours; no volumes were censored or removed from the series. Mean FD was computed from the fMRIPrep confound table, and participants with mean FD > 0.5 mm were excluded to limit motion artefact.

No nuisance regression was applied to the BOLD time series. In particular, the global signal and the mean white-matter and cerebrospinal-fluid signals were deliberately not regressed out: these regressors are derived from — or substantially overlap with — the white-matter BOLD signal that is the object of measurement in this study, and removing them would attenuate or distort the quantity of interest [29]. Head motion was instead addressed at three levels: exclusion of participants with mean FD > 0.5 mm, interpolation of isolated high-motion volumes, and inclusion of mean FD as a covariate in every group-level model. Cardiac and respiratory recordings were not available in the ADNI derivatives used here, so no retrospective physiological correction was possible. Because fALFF is a ratio of band-limited power to total power computed within the same time series, it is invariant to multiplicative scaling of the signal and is comparatively insensitive to between-subject differences in overall signal amplitude [15].

Hippocampal volume and entorhinal cortical thickness were obtained from the ADNI UCSFFSX6 table, which contains outputs of the UCSF cross-sectional FreeSurfer v6 pipeline [30] applied centrally across all ADNI sites. Hippocampal volume was computed as the sum of bilateral hippocampal volumes (fields ST29SV and ST88SV) normalized by intracranial volume (ST10CV). Entorhinal cortical thickness was the mean of bilateral entorhinal thickness estimates (ST24TA and ST83TA).

### 2.3 White Matter Fractional Amplitude of Low-Frequency Fluctuations

The primary imaging predictor was the global white matter fractional amplitude of low-frequency fluctuations (fALFF) in the slow-4 frequency band (0.027–0.073 Hz), a relative measure of spontaneous low-frequency signal amplitude [15]. White matter regions were defined using the Johns Hopkins University (JHU) white-matter tractography atlas, maximum-probability parcellation thresholded at 25% (20 major fiber tracts; JHU-20) [31], as distributed with FSL [32]. The atlas was resampled to the BOLD grid by nearest-neighbour interpolation where required; because atlas and functional data were already defined on the same 2 mm MNI grid, no resampling was performed in practice. Tract labels were applied as provided and were not further restricted by a subject-level tissue-probability mask (see Section 4.5). For each tract, the mean time series across all labelled voxels was demeaned and scaled to unit variance, and its power spectrum was estimated by an unwindowed discrete Fourier transform. No temporal band-pass filter was applied; frequency-band selection was performed in the frequency domain when integrating spectral power. fALFF was computed as the power integrated over the slow-4 band divided by the power integrated from 0.01 Hz to the Nyquist frequency, using trapezoidal integration; because fALFF is a within-series power ratio, it is unaffected by the preceding variance normalization. This implementation differs from the original fALFF definition [15] in two respects: spectral power rather than amplitude (the square root of power) was integrated, and the denominator spanned 0.01 Hz to the Nyquist frequency rather than the full detectable frequency range. These choices were applied consistently across participants; however, because both the power-versus-amplitude choice and the lower bound of the denominator interact with individual differences in spectral shape, this implementation is not equivalent to conventional amplitude-based fALFF and may yield different between-participant associations. Tracts containing no labelled voxels were treated as missing. The global summary measure (G_fALFF_jhu20_slow4) was the unweighted mean across the 20 tract values, so that each tract contributes equally regardless of its size, and served as the primary predictor in all regression models. Prior to analysis, G_fALFF_jhu20_slow4 was z-scored across participants.

The slow-4 band was selected a priori based on three converging lines of evidence. First, slow-4 fALFF demonstrates superior test-retest reliability compared with full-band ALFF and the adjacent slow-5 band (0.01–0.027 Hz), as systematically established by Zuo et al. [16]. Second, Wang et al. [17] demonstrated that slow-4, relative to slow-5, shows broader and more sensitive discrimination of preclinical Alzheimer’s disease stages (SCD, non-amnestic MCI, amnestic MCI) using frequency-specific ALFF. This frequency dependence was first established in amnestic MCI by Han et al. [18], albeit for gray matter and whole-brain rather than white matter BOLD. Third, and most directly relevant to white matter, Wu et al. [19] mathematically characterized a physiologically distinct spectral peak centered at approximately 0.06 Hz in white matter BOLD signals — falling squarely within the slow-4 band — whose amplitude undergoes systematic, age-related attenuation beginning after age 60 and is linked to glymphatic and perivascular clearance mechanisms. Together, these findings indicate that the slow-4 band captures a biologically organized oscillatory component specific to white matter physiology, making it the most appropriate frequency window for WM fALFF quantification.

### 2.4 Structural Imaging Biomarkers

Two classes of structural imaging biomarkers were included as comparison markers. DTI-derived metrics — fractional anisotropy (FA) and mean diffusivity (MD) — were obtained from the ADNI DTIROI_MEAN table (ADNIMERGE2 R package) [23,24,25], which provides regionally averaged DTI metrics computed through ADNI’s centralized DTI processing pipeline. Global white matter FA and MD were computed as the mean across available WM ROIs. Gray matter structural markers — bilateral hippocampal volume and entorhinal cortical thickness — were also obtained from ADNI central processing outputs (UCSFFSX6; see Section 2.2). All four structural markers were matched to the fMRI scan date using the nearest available assessment within a ±365-day window.

### 2.5 Blood-Based Alzheimer’s Disease Biomarkers

Plasma AD biomarkers were obtained from the ADNI UPenn plasma biomarker data release (UPENN_PLASMA_FUJIREBIO_QUANTERIX, March 2026). Four markers were analyzed: phosphorylated tau at threonine-217 (pTau217) and the amyloid-β42/40 ratio (Aβ42/40), both measured using the Fujirebio Lumipulse platform [5,33]; and neurofilament light chain (NfL) and glial fibrillary acidic protein (GFAP), both measured using the Quanterix Simoa platform [4,6]. All four plasma analytes were natural-log transformed before analysis to correct right skew.

Plasma samples were temporally matched to the fMRI scan date using a nearest-date strategy within a ±365-day window. The median gap between plasma collection and fMRI was 38 days (range 0–365 days; 88.7% within 180 days). The complete-case subsample with all four plasma biomarkers and all four structural imaging markers available comprised n = 108 participants.

### 2.6 PET Imaging

Three PET modalities were analyzed. Amyloid PET was quantified using Centiloid units [34] derived from the UC Berkeley Amyloid PET processing pipeline (UCBERKELEY_AMY_6MM) [35]. Tau PET was quantified as the meta-temporal standardized uptake value ratio (META_TEMPORAL_SUVR) from the UC Berkeley Tau PET pipeline (UCBERKELEY_TAU_6MM) [36]. FDG-PET was quantified as the AD-signature region of interest SUVR (SROI.AD) from the BAIPETNMRC dataset; higher SROI.AD values indicate greater glucose metabolism. PET scan dates were matched to fMRI using the same nearest-date strategy as plasma (±365-day window). Each PET modality was analyzed in a separate complete-case sample. All PET outcomes were z-scored prior to analysis.

### 2.7 Cognitive Assessments

Four cognitive outcomes were analyzed. The ADNI episodic memory composite score (ADNI-MEM) [37] and executive function composite score (ADNI-EF) [38] were derived by factor analysis of relevant neuropsychological tests using established ADNI methods. The modified Preclinical Alzheimer’s Cognitive Composite (mPACC) [39] combines tests of memory, orientation, and global cognition and is sensitive to subtle pre-dementia cognitive decline; the Trail Making Test B variant (mPACCtrailsB), in which Trail Making Test Part B replaces the Digit Symbol Substitution Test, was used throughout. The Mini-Mental State Examination (MMSE) [40] served as a global cognition measure. All cognitive scores were z-scored prior to analysis.

### 2.8 Statistical Analyses

All cross-sectional associations were estimated using ordinary least squares (OLS) multiple regression. All continuous predictors were z-scored, and all outcomes were z-scored for the head-to-head analyses to allow direct comparison of standardized effect sizes. Plasma biomarker concentrations were natural-log transformed prior to z-scoring; PET and cognitive outcomes were z-scored without transformation. All models included the following covariates: age, biological sex, years of education, APOE ε4 carrier status, imaging acquisition site (modeled as dummy variables; sites with fewer than five participants were collapsed into an’Other’ category), and mean FD.

Two complementary analytical approaches were applied for each outcome domain. (1) Same-sample single-marker benchmarking: each of the five imaging markers (WM slow-4 fALFF, hippocampal volume, EC thickness, FA, MD) was evaluated in a separate model — marker + covariates — within a common complete-case sample with all five markers and the relevant outcome available. This single-marker-per-model design ensures that partial RZ values reflect each marker’s covariate-adjusted association rather than its incremental contribution beyond competing imaging markers; comparisons across models are valid because all models use the same sample and covariates. For these models we report partial RZ — the proportion of outcome variance left unexplained by the covariate-only model that is explained by adding the imaging marker. (2) Joint-marker modelling with variance decomposition: three model configurations, referred to below as trimmed models, were fitted — DTI-only (WM fALFF + FA + MD), FS-only (WM fALFF + hippocampal volume + EC thickness), and the full model (all four structural markers + WM fALFF). Because the DTI-only and FS-only configurations require only their own markers to be available, they could be fitted in larger, modality-specific complete-case samples than the single-marker benchmark and therefore with greater power; the full model requires all five markers and is fitted in the same complete-case sample as the benchmark. Within each configuration, a commonality analysis partitioned explained variance above the covariate baseline into components unique to fALFF, unique to the structural comparison markers, and a shared (commonality) component. These components are expressed as proportions of total outcome variance, so that they sum to the joint increment over the covariate baseline; they therefore have a different denominator from the partial RZ values reported under (1) and are not directly comparable.

Formally, unique variance attributable to fALFF was defined as RZ_full − RZ_MRI-only (model with MRI markers + covariates, no fALFF); unique variance attributable to MRI was defined as RZ_full − RZ_fALFF-only (model with fALFF + covariates, no MRI); and the commonality coefficient was RZ_MRI + RZ_fALFF − RZ_full − RZ_covariates. Negative commonality values reflect suppression effects and do not imply negative shared variance in a strict set-theoretic sense.

Variance inflation factors (VIF) were computed for all imaging predictors in the full trimmed models. All imaging markers had VIF < 3, indicating acceptable multicollinearity. Statistical significance was set at α = 0.05 (two-tailed). These analyses were hypothesis-driven but not preregistered; results are reported without correction for multiple comparisons as an exploratory characterization of effect magnitudes. FDG-PET analyses (n = 66) are explicitly labeled as secondary exploratory. All analyses were performed in Python 3.13 using NumPy, pandas, and statsmodels.

## 3. Results

### 3.1 Sample Characteristics and Multimodal Data Availability

After resting-state fMRI quality control (mean framewise displacement [FD] ≤ 0.5 mm), 369 ADNI participants with a baseline fMRI session, available age, and available years of education were included. Structural MRI was co-acquired on the same day in 319 participants (86.4%; median gap = 0 days, 100% within 30 days). DTI was available for 287 participants (77.8%; median gap = 0 days, 99.3% within 30 days; range 0–341 days). Plasma AD biomarkers were available in 151 participants (40.9%; median gap = 38 days, 88.7% within 180 days; maximum 365 days). PET scan dates were matched to fMRI using the same ±365-day nearest-date window; the number of participants with each PET measure available is given in Table 1. The complete-case samples for the same-sample PET comparisons, which additionally required all five imaging markers, comprised 207 participants (amyloid Centiloid), 195 (tau meta-temporal SUVR), and 66 (FDG SROI.AD). Demographic and clinical characteristics are reported in Table 1. Multicollinearity among imaging markers was acceptable in all models (VIF < 3 for all predictors in the full model). Head-to-head comparisons of partial RZ across imaging markers are summarized in Figure 1 for the six primary outcomes and in Supplementary Figure 1 for all 11 pre-specified outcomes.

**Figure 1.**
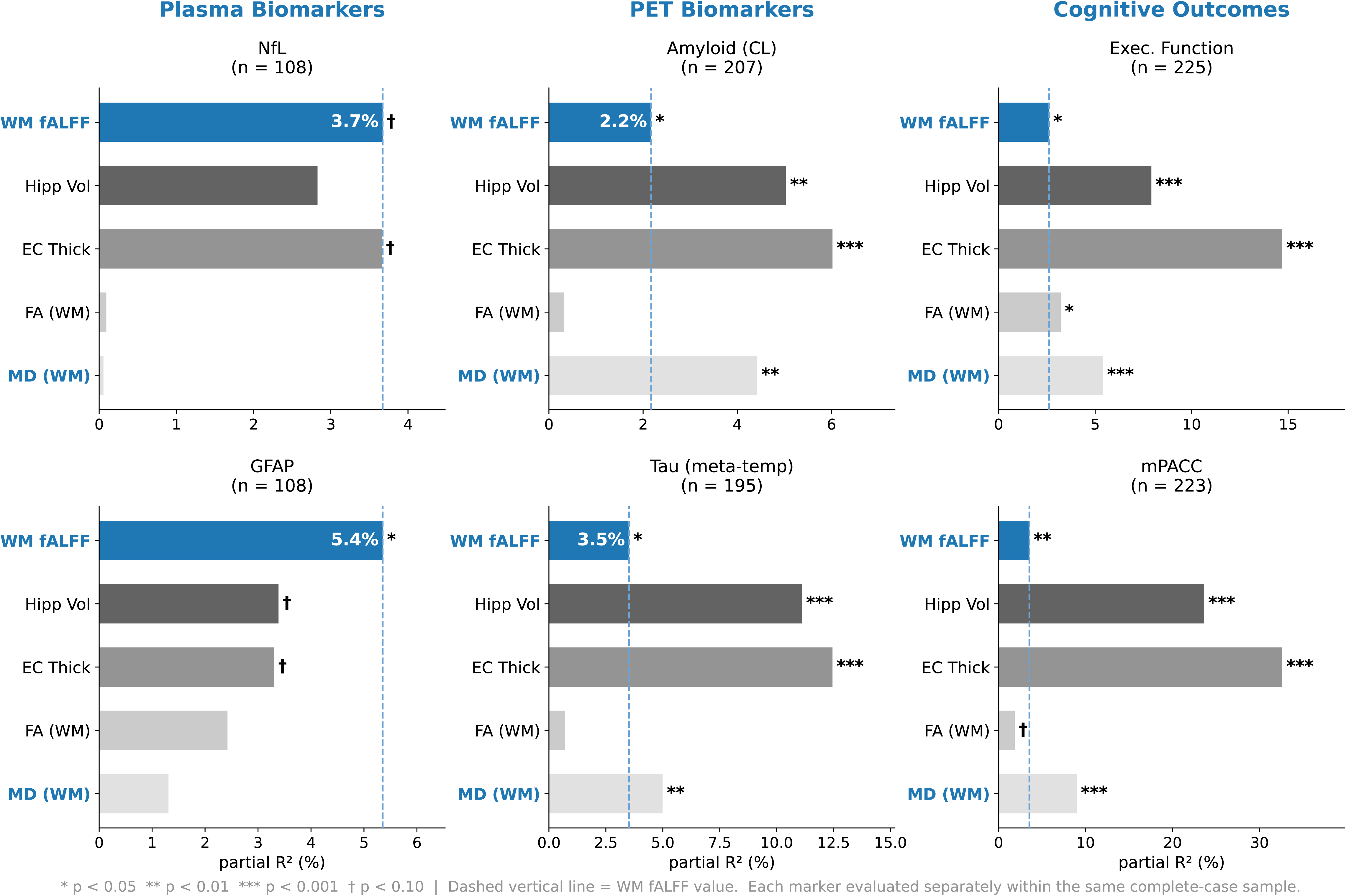
WM slow-4 fALFF compared with conventional imaging markers across the six primary outcomes. Bars show partial RZ (%) for each imaging marker: the proportion of outcome variance left unexplained by the covariate-only model (age, sex, education, APOE ε4 status, site, mean framewise displacement) that is explained by adding that marker. Each marker was evaluated in a separate model within the same complete-case sample for that outcome. Left column, plasma NfL and GFAP (n = 108); middle column, amyloid PET (Centiloid; n = 207) and meta-temporal tau PET SUVR (n = 195); right column, ADNI-EF (n = 225) and mPACC (n = 223). The dashed vertical line marks the WM slow-4 fALFF value for reference. *p < 0.05; **p < 0.01; ***p < 0.001; †p < 0.10. Complete numerical results are given in Tables 2, 4, and 5.

**Table 2.** Same-sample head-to-head comparison of imaging markers for plasma AD biomarkers (n = 108).

| Imaging Marker | pTau217 | Aβ42/40 | NfL | GFAP |
| --- | --- | --- | --- | --- |
|  | β (SE) / partial R<sup>2</sup> | β (SE) / partial R<sup>2</sup> | β (SE) / partial R<sup>2</sup> | β (SE) / partial R<sup>2</sup> |
| WM slow-4 fALFF | **-0.12 (0.10)**<br>partial R<sup>2</sup> = 1.63% | **+0.12 (0.10)**<br>partial R<sup>2</sup> = 1.40% | **-0.17 (0.09)†**<br>partial R<sup>2</sup> = 3.67% | **-0.21 (0.09)\***<br>partial R<sup>2</sup> = 5.35% |
| Hippocampal volume | -0.21 (0.10)*<br>partial R<sup>2</sup> = 4.53% | +0.14 (0.11)<br>partial R<sup>2</sup> = 1.79% | -0.16 (0.10)<br>partial R<sup>2</sup> = 2.83% | -0.18 (0.10)†<br>partial R<sup>2</sup> = 3.39% |
| EC thickness | -0.30 (0.09)**<br>partial R<sup>2</sup> = 10.75% | +0.14 (0.10)<br>partial R<sup>2</sup> = 1.90% | -0.17 (0.09)†<br>partial R<sup>2</sup> = 3.66% | -0.16 (0.09)†<br>partial R<sup>2</sup> = 3.30% |
| FA (WM) | -0.20 (0.11)†<br>partial R<sup>2</sup> = 3.70% | +0.17 (0.11)<br>partial R<sup>2</sup> = 2.51% | -0.03 (0.10)<br>partial R<sup>2</sup> = 0.09% | -0.15 (0.10)<br>partial R<sup>2</sup> = 2.42% |
| MD (WM) | +0.21 (0.10)*<br>partial R<sup>2</sup> = 4.52% | -0.07 (0.11)<br>partial R<sup>2</sup> = 0.38% | +0.02 (0.10)<br>partial R<sup>2</sup> = 0.06% | +0.11 (0.10)<br>partial R<sup>2</sup> = 1.31% |

### 3.2 WM Slow-4 fALFF is Associated with Blood-Based Neurodegeneration Markers

To benchmark WM slow-4 fALFF against conventional imaging markers on equal footing, we evaluated each marker separately within the same complete-case sample (n = 108; all five imaging markers and all four plasma biomarkers available). Each model took the form: plasma ∼ imaging marker + covariates. WM slow-4 fALFF showed a significant covariate-adjusted association with GFAP (β = −0.207, SE = 0.092, p = 0.026, partial RZ = 5.35%), with higher fALFF associated with lower GFAP. This partial RZ was comparable to or larger than that of hippocampal volume (3.39%), EC thickness (3.30%), FA (2.42%), and MD (1.31%) in the same sample. No significant covariate-adjusted associations were observed for fALFF with NfL, pTau217, or Aβ42/40. Full same-sample benchmarking results are reported in Table 2 and Figure 1 (left column).

To assess the incremental value of WM fALFF beyond conventional DTI white matter markers, we fitted modality-specific trimmed models. In the DTI-only trimmed model (fALFF + FA + MD; no gray matter markers; n = 121), fALFF showed significant unique variance for both NfL (β = −0.223, p = 0.010; unique fALFF = 3.65%) and GFAP (β = −0.247, p = 0.004; unique fALFF = 4.45%). Notably, fALFF’s unique contribution to NfL (3.65%) was nearly 8-fold larger than DTI’s combined unique contribution (0.44%), demonstrating that functional white matter activity captures neurodegeneration-related variance well beyond what is explained by FA and MD. In the FS-only trimmed model (fALFF + hippocampal volume + EC thickness; n = 134), fALFF remained significantly associated with NfL (β = −0.176, p = 0.046; unique = 2.11%) and GFAP (β = −0.206, p = 0.021; unique = 2.89%), exceeding the combined unique contribution of the gray matter markers (hippocampal volume + EC thickness: NfL 1.71%, GFAP 0.49%). When all five imaging markers were entered simultaneously (n = 108), fALFF was no longer significantly associated with either NfL (β = −0.135, p = 0.173) or GFAP (β = −0.155, p = 0.117), indicating substantial shared variance with the structural markers. Trimmed model variance decomposition results are reported in Table 3.

**Table 3.**
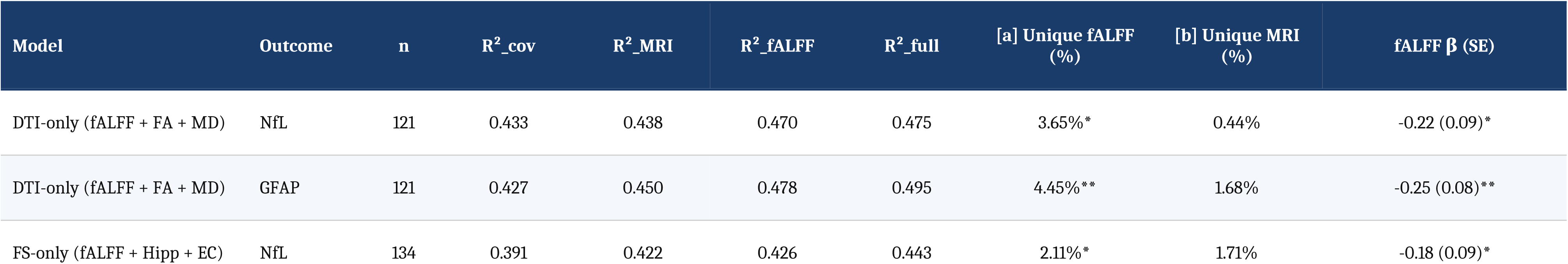
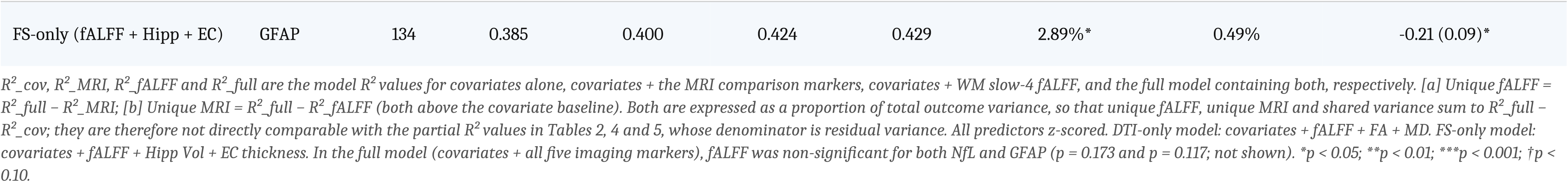
Trimmed model variance decomposition for plasma NfL and GFAP (DTI-only and FS-only models).

**Table 4.**
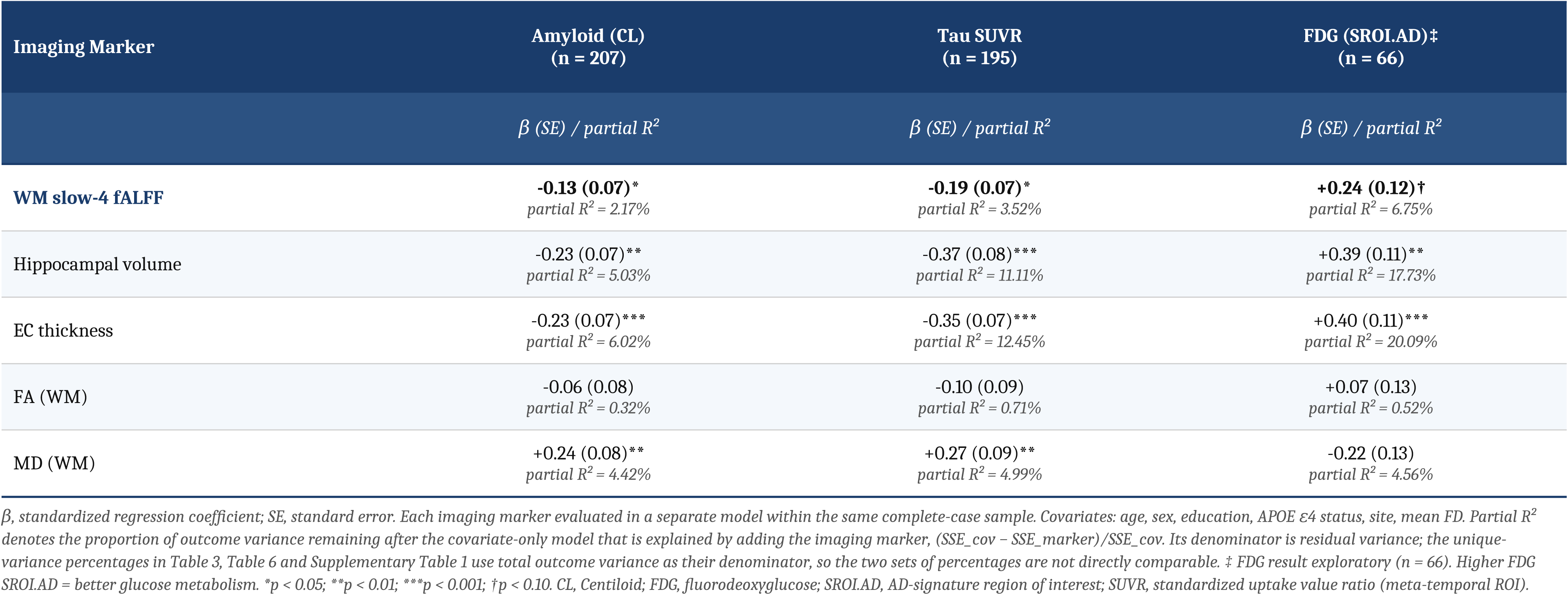
Same-sample head-to-head comparison of imaging markers for PET biomarkers.

### 3.3 WM Slow-4 fALFF is Associated with Amyloid and Tau PET

Using the same benchmarking approach — each imaging marker evaluated separately within a common complete-case sample (PET + all five imaging markers + covariates) — WM slow-4 fALFF showed significant covariate-adjusted associations with amyloid burden (Centiloid; β = −0.134, SE = 0.067, p = 0.046, partial RZ = 2.17%; n = 207) and tau burden (meta-temporal SUVR; β = −0.187, SE = 0.075, p = 0.013, partial RZ = 3.52%; n = 195), with higher fALFF associated with lower pathological burden. Gray matter markers showed larger partial RZ for PET pathology in the same sample (hippocampal volume: 5.03% amyloid, 11.11% tau; EC thickness: 6.02% amyloid, 12.45% tau), as did MD (amyloid: p = 0.004; tau: p = 0.003). When all five imaging markers were entered simultaneously, fALFF associations with PET were attenuated to non-significance (amyloid: β = −0.088, p = 0.182; tau: β = −0.098, p = 0.172), reflecting shared variance with gray matter markers. Full same-sample benchmarking results are reported in Table 4 and Figure 1 (middle column).

In the DTI-only PET trimmed model (*n* = 237), fALFF showed significant incremental value for amyloid burden independent of FA and MD (β = −0.153, *p* = 0.012; unique fALFF = 1.94%; unique DTI = 2.57%). Trimmed model variance decomposition results are reported in Supplementary Table 1.

### 3.4 WM Slow-4 fALFF is Associated with Executive Function and Composite Cognition

Using the same benchmarking approach — each imaging marker evaluated separately within a common complete-case sample — WM slow-4 fALFF showed significant covariate-adjusted associations with executive function (ADNI-EF; β = +0.149, SE = 0.064, p = 0.022, partial RZ = 2.61%; n = 225) and the mPACC composite score (β = +0.171, SE = 0.064, p = 0.008, partial RZ = 3.53%; n = 223), with higher fALFF associated with better performance. Gray matter markers showed substantially larger partial RZ for cognition in the same sample (EC thickness: 14.7–32.6%; hippocampal volume: 7.9–23.6%), and DTI markers were also significant (FA: EF p = 0.011; MD: EF and mPACC both p < 0.001). When all five imaging markers were entered simultaneously, fALFF associations with cognition were attenuated to non-significance (EF: β = +0.069, p = 0.258; mPACC: β = +0.051, p = 0.326), reflecting shared variance with gray matter and DTI markers. Full same-sample benchmarking results are reported in Table 5 and Figure 1 (right column).

**Table 5.** Same-sample head-to-head comparison of imaging markers for cognitive outcomes.

| Imaging Marker | ADNI-MEM<br>(n = 225) | ADNI-EF<br>(n = 225) | mPACC<br>(n = 223) | MMSE<br>(n = 222) |
| --- | --- | --- | --- | --- |
| | $\beta$ (SE) / partial $R^2$ | $\beta$ (SE) / partial $R^2$ | $\beta$ (SE) / partial $R^2$ | $\beta$ (SE) / partial $R^2$ |
| **WM slow-4 fALFF** | **+0.11 (0.06)†**<br>partial $R^2$ = 1.68% | **+0.15 (0.06)\***<br>partial $R^2$ = 2.61% | **+0.17 (0.06)\*\***<br>partial $R^2$ = 3.53% | **+0.11 (0.07)†**<br>partial $R^2$ = 1.43% |
| Hippocampal volume | +0.41 (0.07)***<br>partial $R^2$ = 16.44% | +0.30 (0.07)***<br>partial $R^2$ = 7.91% | +0.51 (0.06)***<br>partial $R^2$ = 23.62% | +0.48 (0.07)***<br>partial $R^2$ = 18.93% |
| EC thickness | +0.50 (0.06)***<br>partial $R^2$ = 29.24% | +0.37 (0.06)***<br>partial $R^2$ = 14.69% | +0.55 (0.06)***<br>partial $R^2$ = 32.60% | +0.51 (0.06)***<br>partial $R^2$ = 26.06% |
| FA (WM) | +0.13 (0.08)†<br>partial $R^2$ = 1.37% | +0.21 (0.08)*<br>partial $R^2$ = 3.21% | +0.16 (0.08)†<br>partial $R^2$ = 1.84% | +0.19 (0.09)*<br>partial $R^2$ = 2.38% |
| MD (WM) | -0.29 (0.08)***<br>partial $R^2$ = 6.45% | -0.27 (0.08)***<br>partial $R^2$ = 5.40% | -0.35 (0.08)***<br>partial $R^2$ = 8.97% | -0.31 (0.08)***<br>partial $R^2$ = 6.25% |

In the DTI-only cognitive trimmed model (fALFF + FA + MD; *n* = 253–255), fALFF explained significant incremental variance in EF (β = +0.117, *p* = 0.048; unique fALFF = 1.13%) and mPACC (β = +0.161, *p* = 0.006; unique fALFF = 2.14%) beyond FA and MD. In contrast to the plasma analyses, DTI’s unique cognitive contribution (3.7–4.9%) exceeded fALFF’s in this context, suggesting that WM fALFF preferentially captures neurodegeneration/neuroinflammatory burden rather than purely cognitive variance. Cognitive trimmed model variance decomposition results are reported in Table 6.

**Table 6.** Trimmed model variance decomposition for cognitive outcomes (DTI-only model).

| Outcome | n | $R^2_{cov}$ | $R^2_{MRI}$ | $R^2_{fALFF}$ | $R^2_{full}$ | [a] Unique fALFF (%) | [b] Unique DTI (%) | fALFF $\beta$ (SE) |
| --- | --- | --- | --- | --- | --- | --- | --- | --- |
| ADNI-MEM | 255 | 0.359 | 0.402 | 0.370 | 0.410 | 0.79%† | 3.99% | +0.10 (0.06)† |
| ADNI-EF | 255 | 0.298 | 0.340 | 0.314 | 0.351 | 1.13%* | 3.72% | +0.12 (0.06)* |
| mPACC | 254 | 0.297 | 0.351 | 0.324 | 0.373 | 2.14%** | 4.88% | +0.16 (0.06)** |
| MMSE | 253 | 0.226 | 0.270 | 0.240 | 0.280 | 1.04%† | 4.02% | +0.11 (0.06)† |

DTI-only model: covariates + WM slow-4 fALFF + FA (WM) + MD (WM); no gray matter markers. FS-only and full models: fALFF was null for all cognitive outcomes (all $p > 0.20$ ; not shown). $R^2_{cov}$ , $R^2_{MRI}$ , $R^2_{fALFF}$ and $R^2_{full}$ are the model $R^2$ values for covariates alone, covariates + FA and MD, covariates + WM slow-4 fALFF, and the full DTI-only model containing both, respectively. [a] Unique fALFF = $R^2_{full} - R^2_{MRI}$ ; [b] Unique DTI = $R^2_{full} - R^2_{fALFF}$ (both above the covariate baseline). Both are expressed as a proportion of total outcome variance, so that unique fALFF, unique DTI and shared variance sum to $R^2_{full} - R^2_{cov}$ ; they are therefore not directly comparable with the partial $R^2$ values in Tables 2, 4 and 5, whose denominator is residual variance. Covariates: age, sex, education, APOE $\epsilon 4$ status, site, mean FD. \* $p < 0.05$ ; \*\* $p < 0.01$ ; † $p < 0.10$ . ADNI-MEM, ADNI-EF = ADNI memory and executive function composites; mPACC, modified Preclinical Alzheimer's Cognitive Composite (Trail Making Test B variant, mPACCtrailsB).

## 4. Discussion

The present study systematically benchmarked global white matter (WM) slow-4 fractional amplitude of low-frequency fluctuations (fALFF) against four established structural imaging markers — hippocampal volume, entorhinal cortical (EC) thickness, fractional anisotropy (FA), and mean diffusivity (MD) — across 11 pre-specified outcome domains in ADNI. WM slow-4 fALFF showed significant covariate-adjusted associations with plasma GFAP and, in larger modality-specific samples, with NfL, amyloid and tau PET, executive function, and the mPACC composite. Variance decomposition revealed that functional WM activity explains unique neurodegeneration-related variance beyond DTI microstructure.

### 4.1 WM fALFF and Plasma Neurodegeneration Markers

Among the plasma biomarkers examined, the association between WM slow-4 fALFF and glial fibrillary acidic protein (GFAP) was the most prominent finding, with fALFF explaining a larger proportion of the residual GFAP variance remaining after covariate adjustment (partial RZ = 5.35%) than hippocampal volume (3.39%), EC thickness (3.30%), FA (2.42%), or MD (1.31%) in the same complete-case sample. GFAP is a marker of astroglial reactivity and is classified as an Inflammatory/Immune (I-class) biomarker under the revised 2024 Alzheimer’s Association diagnostic framework [4]. Pereira et al. demonstrated that elevated plasma GFAP reflects amyloid-β-driven astroglial activation and is detectable even in the preclinical stage, prior to overt tau pathology or neurodegeneration [41]. The magnitude of the fALFF-GFAP association, and its relative advantage over structural markers, suggests that low-frequency WM functional dynamics may be sensitive to the same pathological milieu of astroglial reactivity, a finding consistent with the multi-modal GFAP-imaging associations documented by Shir et al. in a Mayo Clinic cohort [42].

In trimmed variance decomposition models, WM fALFF also provided substantial unique variance for NfL beyond FA and MD (unique ΔRZ = 3.65%), nearly eight times the unique contribution of DTI (0.44%). NfL is an established N-class marker of neuroaxonal injury and is non-specific to AD etiology [4,6]. Nabizadeh et al. demonstrated using ADNI data that plasma NfL correlates with white matter microstructural damage quantified by DTI [10], and Benedet et al. showed that NfL-imaging associations are disease-stage-specific, with stronger links to tau and cortical thinning at later stages [43]. In amyloid-positive individuals, plasma NfL has also been linked to AD-related metabolic decline on FDG-PET [44]. The disproportionately large unique contribution of fALFF to NfL variance, relative to FA and MD, suggests that functional white matter signal dynamics capture neuroaxonal injury-related information that is not fully reflected in diffusion-based microstructural indices. This may reflect the sensitivity of resting-state BOLD fluctuations to integrated axonal and glial activity across entire WM fiber systems, extending beyond local fiber organization measured by DTI. This advantage held against each structural modality separately — fALFF also explained significant unique variance beyond hippocampal volume and EC thickness (NfL, p = 0.046; GFAP, p = 0.021) — but not against both together: in the full five-marker model (n = 108), the unique contribution of fALFF to NfL and GFAP was no longer significant (p = 0.173 and p = 0.117), indicating that much of the variance fALFF captures is shared with the structural markers collectively. The fALFF–NfL association was not significant in the single-marker benchmark sample (n = 108, p = 0.067) but was significant in the DTI-only model (n = 121, p = 0.010); this difference may reflect changes in sample size, sample composition and model specification, which varied together across the two analyses.

### 4.2 WM fALFF and PET Biomarkers

WM slow-4 fALFF was significantly associated with both amyloid burden (Centiloid; partial RZ = 2.17%) and tau burden (meta-temporal SUVR; partial RZ = 3.52%), with higher fALFF corresponding to lower pathological burden. These effect sizes were smaller than those of gray matter markers in the same samples, particularly EC thickness (amyloid: 6.02%; tau: 12.45%) and hippocampal volume (amyloid: 5.03%; tau: 11.11%), and fALFF associations were attenuated to non-significance when all five imaging markers were modeled simultaneously, indicating substantial shared variance with established structural biomarkers. Gonzales et al. similarly observed that WM DTI microstructure is correlated with both amyloid and tau PET in ADNI, with different association patterns across PET modalities [9]. The present findings extend this by showing that slow-4 fALFF carries sensitivity to amyloid and tau pathology comparable to that of DTI microstructure on a per-marker basis while explaining partly non-overlapping variance (incremental fALFF contribution for amyloid: 1.94% beyond FA and MD in the DTI-only trimmed model), supporting a complementary rather than redundant relationship between functional and structural WM imaging.

### 4.3 WM fALFF and Cognitive Outcomes

WM slow-4 fALFF was significantly associated with executive function (ADNI-EF; partial RZ = 2.61%) and the mPACC composite (partial RZ = 3.53%), but not with episodic memory or MMSE. This dissociation is notable: executive function and the mPACC, which weights orientation and global cognition, may engage WM networks more directly than purely hippocampal-dependent memory measures. In the DTI-only trimmed models, fALFF provided significant unique variance for EF and mPACC (1.13% and 2.14%, respectively), whereas DTI’s unique cognitive contribution (3.7–4.9%) was proportionally larger than for the plasma biomarkers. Gray matter structural markers showed substantially larger cognitive associations (EC thickness: 14.7–32.6%; hippocampal volume: 7.9–23.6%), and fALFF was attenuated to non-significance when combined with all markers simultaneously. Together, these patterns suggest that WM functional activity preferentially captures neurodegeneration-and neuroinflammation-related variance — as reflected in plasma NfL and GFAP — rather than serving as a primary driver of cognitive differences, which are dominated by gray matter atrophy in this ADNI sample spanning CN to AD dementia.

### 4.4 Physiological and Biological Interpretation

The biological basis of slow-4 WM fALFF likely reflects multiple interacting contributors. The fALFF metric quantifies the relative power of spontaneous low-frequency BOLD fluctuations within a specific frequency band and, when applied to WM voxels, may integrate contributions from glial metabolic activity, perivascular and glymphatic dynamics, axonal conduction-coupled hemodynamics, and vessel wall compliance [29]. Intracranial electrophysiological evidence from SEEG recordings has demonstrated that WM BOLD functional connectivity co-varies with electrophysiological synchrony across multiple frequency bands [14], lending neural validity to WM functional signals. The specificity of the slow-4 band (0.027–0.073 Hz) is further supported by the observation of a distinct physiological spectral peak in WM BOLD at approximately 0.06 Hz — within the slow-4 range — that shows age-related attenuation and is linked to glymphatic and perivascular mechanisms [19]. The preferential association of slow-4 fALFF with astroglial (GFAP) and neuroaxonal injury (NfL) markers, relative to its more modest associations with amyloid and tau pathology per se, is consistent with the interpretation that WM functional disruption may represent a downstream consequence of integrated white matter pathology — including demyelination, axonal loss, and reactive gliosis — rather than a specific marker of amyloid or tau aggregation. We recommend describing WM fALFF as reflecting low-frequency BOLD signal dynamics, and note that direct equivalence to neuronal action potentials should not be inferred.

### 4.5 Strengths, Limitations, and Future Directions

Strengths of this study include the use of a large, well-characterized multimodal dataset (ADNI), a systematic head-to-head benchmarking design within matched complete-case samples ensuring fair comparisons, and variance decomposition analyses that quantify unique versus shared contributions. The a priori selection of the slow-4 frequency band, grounded in convergent evidence regarding reliability, AD sensitivity, and WM-specific spectral properties, mitigates concerns about selective frequency reporting.

Several limitations warrant acknowledgment. The cross-sectional design precludes causal inference and limits assessment of whether WM fALFF predicts longitudinal disease trajectory. Complete-case analyses, particularly for plasma biomarkers (n = 108) and FDG-PET (n = 66), reduce statistical power and may introduce selection bias. WM fALFF was quantified from atlas-defined tract labels without intersecting a subject-level white-matter probability map, and no partial-volume correction was applied; at 2 mm resolution, voxels near tract borders may therefore contain gray matter or cerebrospinal fluid, and residual partial-volume contamination cannot be excluded. Relatedly, no nuisance signals were regressed from the BOLD time series (Section 2.2), so unmodelled physiological and scanner-related variance remains in the fALFF estimates, although the fALFF ratio normalizes for multiplicative signal scaling and mean FD was included as a covariate throughout. The ADNI sample spans the full clinical spectrum from cognitively normal to AD dementia, and associations may differ in enriched preclinical cohorts. These analyses were hypothesis-driven but not formally preregistered; results are reported without correction for multiple comparisons and should be considered exploratory characterizations of effect magnitudes. The physiological interpretation of WM BOLD signals remains an active area of investigation, and caution is warranted in attributing fALFF changes to specific cellular mechanisms. Future work should examine slow-4 fALFF at the tract level, integrate longitudinal designs, and test its sensitivity in amyloid-positive cognitively normal individuals — the population most relevant to secondary prevention trials.

### 4.6 Conclusion

WM slow-4 fALFF captures neurodegeneration-related variance — particularly astroglial reactivity and neuroaxonal injury as indexed by plasma GFAP and NfL — that is incremental to DTI white matter microstructure and partially independent of gray matter atrophy. Associations with amyloid and tau PET and with cognitive composites are present but reflect substantial shared variance with established structural markers. These findings position functional WM imaging, specifically slow-4 band fALFF over major WM tracts, as a complementary non-invasive marker within the multimodal AD biomarker landscape, with potential value for characterizing integrated white matter pathology in aging and AD research.

## Data availability

Data used in the preparation of this article were obtained from the Alzheimer’s Disease Neuroimaging Initiative (ADNI) database (adni.loni.usc.edu). Individual-level ADNI imaging, plasma biomarker, PET and clinical data are available to qualified investigators following application and approval through the ADNI Data Sharing and Publications Committee (https://adni.loni.usc.edu/data-samples/access-data/); owing to ADNI data-access policies, the authors are not permitted to redistribute individual-level data. Participant-level measures derived from these data, including the tract-wise white matter fALFF values, are likewise covered by that restriction and cannot be redistributed. The summary-level results supporting all conclusions are reported in full in Tables 1–6, Supplementary Table 1, Figure 1 and Supplementary Figure 1.

ADNI data tables used in this study: UPENN_PLASMA_FUJIREBIO_QUANTERIX (plasma pTau217, Aβ42/40, NfL, GFAP); UCBERKELEY_AMY_6MM (amyloid PET Centiloid); UCBERKELEY_TAU_6MM (meta-temporal tau SUVR); BAIPETNMRC (FDG SROI.AD); UCSFFSX6 (FreeSurfer v6 hippocampal volume and entorhinal thickness); DTIROI_MEAN (regional FA and MD); and ADNIMERGE2 (demographics, APOE ε4 status, diagnosis and cognitive composites). The JHU white-matter tractography atlas is distributed with FSL.

## Code availability

Custom code for the full analysis pipeline — tract-wise white matter slow-4 fALFF derivation, same-sample head-to-head benchmarking, modality-specific trimmed models, and commonality-based variance decomposition — will be made publicly available upon peer-reviewed publication, and is available from the corresponding author on reasonable request in the interim. All analysis steps are described in the Methods in sufficient detail to permit independent reimplementation from the ADNI tables listed above.

The analyses used the following publicly available software: fMRIPrep 25.1.4 (https://fmriprep.org/); FSL v6.0.5 (https://fsl.fmrib.ox.ac.uk/), which distributes the JHU white-matter tractography atlas; FreeSurfer v6 (https://surfer.nmr.mgh.harvard.edu/), applied centrally by ADNI; and Python 3.13 with NumPy, pandas, nibabel, nilearn and statsmodels.

## Author contributions

Y.H. performed the data processing, statistical analyses and visualization, and wrote the original draft. L.Z. conceived and designed the study and supervised the work. A.W.T. and L.Z. provided funding. All authors reviewed and edited the manuscript and approved the final version.

CRediT roles: Conceptualization, Y.H. and L.Z.; Methodology, Y.H. and L.Z.; Software, Y.H. and L.Z.; Formal analysis, Y.H. and L.Z.; Investigation, Y.H., L.Z. and A.W.T.; Data curation, Y.H. and L.Z.; Writing — original draft, Y.H.; Writing — review & editing, Y.H., A.W.T. and L.Z.; Visualization, Y.H. and L.Z.; Supervision, L.Z.; Funding acquisition, A.W.T. and L.Z.

## Acknowledgements

Data collection and sharing for this project was funded by the Alzheimer’s Disease Neuroimaging Initiative (ADNI) (National Institutes of Health Grant U19 AG024904) and DOD ADNI (Department of Defense award number W81XWH-12-2-0012). ADNI is funded by the National Institute on Aging, the National Institute of Biomedical Imaging and Bioengineering, and through generous contributions from the following: AbbVie, Alzheimer’s Association; Alzheimer’s Drug Discovery Foundation; Araclon Biotech; BioClinica, Inc.; Biogen; Bristol-Myers Squibb Company; CereSpir, Inc.; Cogstate; Eisai Inc.; Elan Pharmaceuticals, Inc.; Eli Lilly and Company; EuroImmun; F. Hoffmann-La Roche Ltd and its affiliated company Genentech, Inc.; Fujirebio; GE Healthcare; IXICO Ltd.; Janssen Alzheimer Immunotherapy Research & Development, LLC.; Johnson & Johnson Pharmaceutical Research & Development LLC.; Lumosity; Lundbeck; Merck & Co., Inc.; Meso Scale Diagnostics, LLC.; NeuroRx Research; Neurotrack Technologies; Novartis Pharmaceuticals Corporation; Pfizer Inc.; Piramal Imaging; Servier; Takeda Pharmaceutical Company; and Transition Therapeutics. The Canadian Institutes of Health Research is providing funds to support ADNI clinical sites in Canada. Private sector contributions are facilitated by the Foundation for the National Institutes of Health (www.fnih.org). The grantee organization is the Northern California Institute for Research and Education, and the study is coordinated by the Alzheimer’s Therapeutic Research Institute at the University of Southern California. ADNI data are disseminated by the Laboratory for Neuro Imaging at the University of Southern California.

This work was additionally supported by the NIH SPAN Coordinating Center (U24NS130600) and by NIH Grant 5R21MH133239-02. Research reported in this publication was supported by the Office of the Director, National Institutes of Health, under Award Number S10OD032285.

## Tables

Tables 1–6 and Supplementary Table 1 are not embedded in this manuscript file; they are supplied as separate files (Table 1: Table_1.docx; Tables 2–6: Tables_2-6.docx; Supplementary Table 1: Supplementary_Table_1.docx), each with its complete footnote.

**Supplementary Table 1. Trimmed model variance decomposition for PET biomarkers (DTI-only model).**

## Figures

Figures are supplied as separate image files (Figure 1: Figure_1.png / Figure_1.pdf; Supplementary Figure 1: Supplementary_Figure_1.png / Supplementary_Figure_1.pdf) and are not embedded in this manuscript file. Legends are given below.

Supplementary Figure 1. WM slow-4 fALFF compared with conventional imaging markers across all 11 pre-specified outcomes. Panels and conventions as in Figure 1, additionally showing plasma pTau217 and Aβ42/40, FDG-PET (SROI.AD; n = 66, exploratory), ADNI-MEM (n = 225), and MMSE (n = 222). The dashed vertical line marks the WM slow-4 fALFF value for reference. Complete numerical results are given in Tables 2, 4, and 5.

